# Schizophrenia-associated *NRXN1* deletions induce developmental-timing- and cell-type-specific vulnerabilities in human brain organoids

**DOI:** 10.1101/2022.08.24.505165

**Authors:** Rebecca Sebastian, Kang Jin, Narciso Pavon, Ruby Bansal, Andrew Potter, Yoonjae Song, Juliana Babu, Rafael Gabriel, Yubing Sun, Bruce Aronow, ChangHui Pak

## Abstract

*De novo* mutations and copy number deletions in *NRXN1* (2p16.3) pose a significant risk for schizophrenia (SCZ). It is unclear how *NRXN1* deletions impact cortical development in a cell type-specific manner and disease background modulates these phenotypes. Here, we leveraged human pluripotent stem cell-derived forebrain organoid models carrying *NRXN1* heterozygous deletions in isogenic and SCZ patient genetic backgrounds and conducted single-cell transcriptomic analysis over the course of brain organoid development from 3 weeks to 3.5 months. Intriguingly, while both deletions similarly impacted molecular pathways associated with ubiquitin-proteasome system, alternative splicing, and synaptic signaling in maturing glutamatergic and GABAergic neurons, SCZ-*NRXN1* deletions specifically perturbed developmental trajectories of early neural progenitors and accumulated disease-specific transcriptomic signatures. Using calcium imaging, we found that both deletions led to long-lasting changes in spontaneous and synchronous neuronal networks, implicating synaptic dysfunction. Our study reveals developmental-timing- and cell-type-dependent actions of *NRXN1* deletions in unique genetic contexts.

---

*De novo* mutations and copy number variations (CNVs) in 2p16.3 have been repeatedly observed in patients with autism spectrum disorders (ASDs), SCZ, and intellectual disability^1–3^. Albeit rare, these CNV losses usually manifest in heterozygous deletions, and present a significant increase in risk for multiple neuropsychiatric disorders^4^. Neurexin-1 (*NRXN1*), the single gene present in this locus, encodes a type I membrane cell adhesion molecule that functions as a synaptic organizer at central synapses^5^. As a presynaptic molecule, NRXN1 associates with multiple soluble and transmembrane molecules, thereby endowing specific synapses with unique synaptic signaling and transmission properties^6–9^. *NRXN1* also undergoes extensive alternative splicing, further enriching the diversity of these interactions^10–13^. Due to this pan-synaptic role throughout the brain, it is not surprising to find strong prevalence of *NRXN1* genetic lesions in multiple neurodevelopmental and psychiatric disorders. Most often, these lesions are large deletions (up to ∼1Mb) affecting the long isoform NRXN1α specifically, as well as *NRXN1α /ý* lesions affecting both the long and short isoforms. However, it remains unknown why or how the same *NRXN1* deletion results in phenotypically distinct disorders in individuals. It is often hypothesized that the interaction between common variants (disease genetic background) and *NRXN1* CNVs drives these differences. Yet, experimentally demonstrating this hypothesis has been challenging.

Separate from its canonical function at synapses which occurs post-neurogenesis, recent evidence suggests possible roles of *NRXN1* in early cortical development. In fact, *NRXN1* mRNAs are abundantly detected in human embryonic neocortex, as early as gestational week (GW) 14, reaching peak at birth before slowly decreasing with age^14^. Knockdown of *NRXN1* in human neural progenitor cells (NPCs) results in decreased levels of glial progenitor marker GFAP, thereby potentially skewing the ratio of neurons to astrocytes^15^. A bi-allelic *NRXN1α* deletion in human iPSC-derived neural cells has been shown to impair maturation of neurons and shift NPC differentiation potential towards glial rather than neuronal fate^16^. More recently, *in vivo* CRISPR KO of *nrxn1* in *Xenopus tropicalis* embryos led to increased telencephalon size attributed to the increased proliferation of NPCs^17^. Separate validation using human cortical NPCs and iPSC-derived organoids showed increased proliferation of NPCs and an increase in neurogenesis in *NRXN1* mutants^17^. Though these studies provide some clues as to which roles *NRXN1* may play during early corticogenesis, the outcomes from these distinct models are inconsistent due to the differences in genetic lesions and the dosage of *NRXN1* being manipulated at different developmental time points. Therefore, it is worth investigating whether disease-associated *NRXN1* mutations in human cells lead to aberrant cortical development and maturation trajectories of neuronal populations in the cortex, thereby ultimately impacting cortical circuitry and synaptic function.

Human pluripotent stem cell (hPSC) derived brain organoids have been proven useful to model early developmental processes associated with neuropsychiatric disorders^18–24^. The self-organizing capability of hPSCs under directly guided differentiation produces relatively homogeneous brain organoids, which can be maintained under defined conditions over long term^20, 25^. By deriving cortical brain organoids from human induced pluripotent stem cells (iPSCs) representing a heterogeneous population of SCZ individuals, studies showed that there exists differentially regulated transcriptomic profiles^26^, neuronal synaptic transmission defects^27^, and early cortical maldevelopment^28^. More recently, brain organoids derived from idiopathic SCZ iPSCs exhibited reduced capacity to differentiate into neurons from NPCs^29^. Though these studies are promising and provide certain clues to the early brain developmental mechanisms of SCZ, it remains unclear how certain cell types during a continuous developmental time window are affected in such human cellular models of SCZ and how specific disease risk variants affect this process. More importantly, since genetic backgrounds often contribute to and modulate cellular phenotypes, understanding even how a single disease variant acts is difficult to dissect unless an isogenic mutant model is analyzed side by side with the patient model.

Here, we differentiated brain organoids from a panel of hPSC lines, where *NRXN1* CNVs have been either artificially engineered (isogenic) or deleted genetically in individuals with SCZ (patient iPSCs) paired with controls^30, 31^. From 3 weeks up to 3.5 months, we collected and generated a total of 156,966 high-quality single cell transcriptomes and performed an in-depth analysis on the neurodevelopmental impact of *NRXN1* deletions in both engineered and SCZ genetic backgrounds. Over brain organoid development, we found that SCZ-*NRXN1* deletions affect developmental trajectories of early neural progenitors as early as 3 weeks, whereas engineered *NRXN1* deletions act at later stages of development influencing neuronal maturation. The degree of transcriptional perturbations in various cell types, as well as the accumulation of disease-specific signatures, were enhanced in the patient deletions compared to engineered deletions. At the molecular level, we identified that maturing glutamatergic and GABAergic neurons are consistently impacted due to *NRXN1* deletions irrespective of genetic background, attributed in part to altered gene expression programs in ubiquitin-proteasome system, alternative splicing, and synaptic signaling. Detailed analysis of differentially expressed genes (DEGs) revealed that neuronal splicing regulators are specifically misregulated, rendering an immature global splicing program. By performing complementary analysis using bulk RNA sequencing dataset from 2D induced neurons, we uncovered differential isoform abundance and local splicing misregulation in *NRXN1* deletions, including altered splicing patterns of *NRXN1* gene itself. At the functional level, calcium imaging analysis of brain organoids showed long-lasting changes in spontaneous and synchronous neuronal networks, implicating synaptic dysfunction across backgrounds. Our study reveals developmental-timing-and cell-type-dependent actions of *NRXN1* deletions in unique genetic contexts and elucidates potential molecular and cellular origins of synaptic dysfunction.

## Generation of forebrain organoids for modeling NRXN1 deletions in neocortical development

We generated dorsal forebrain organoids as previously described, chosen for its reported homogeneity of the cellular constituents and reproducibility in disease modeling^27, 32^ (Fig. 1a). Patterned forebrain organoids showed expected developmental milestones as previously reported. In early time point at day 21, actively dividing proliferative ventricular zones (MKI67, SOX2) appeared, which decreased in abundance over the course of maturation (Fig. S1-2). Starting at day 50 and well into day 100, the spatial organization of HOPX+ outer radial glia (oRGs), EOMES+ intermediate progenitor cells (IPCs), and BCL11B+ deeper layer and SATB2+ upper layer cortical neurons were detected (Fig. S1-2). Moreover, the presence of S100B+ developing astrocytes and NEUN+ mature neurons were reliably detected at day 100 (Fig. S1-2), indicative of active neurogenesis and the start of astrogenesis. At this time point and beyond, presynaptic markers (SYNAPSIN and SYNAPTOPHYSIN) and postsynaptic marker (HOMER) were also detected along MAP2+ dendrites (Fig. S1-2), suggesting that the developing neurons are actively forming synapses in this organoid model. Thus, we chose these three developmental time points for 10X single-cell RNA sequencing (scRNAseq), thereby capturing the different pools of cell identities undergoing fate specification and maturation.

**Figure 1.**
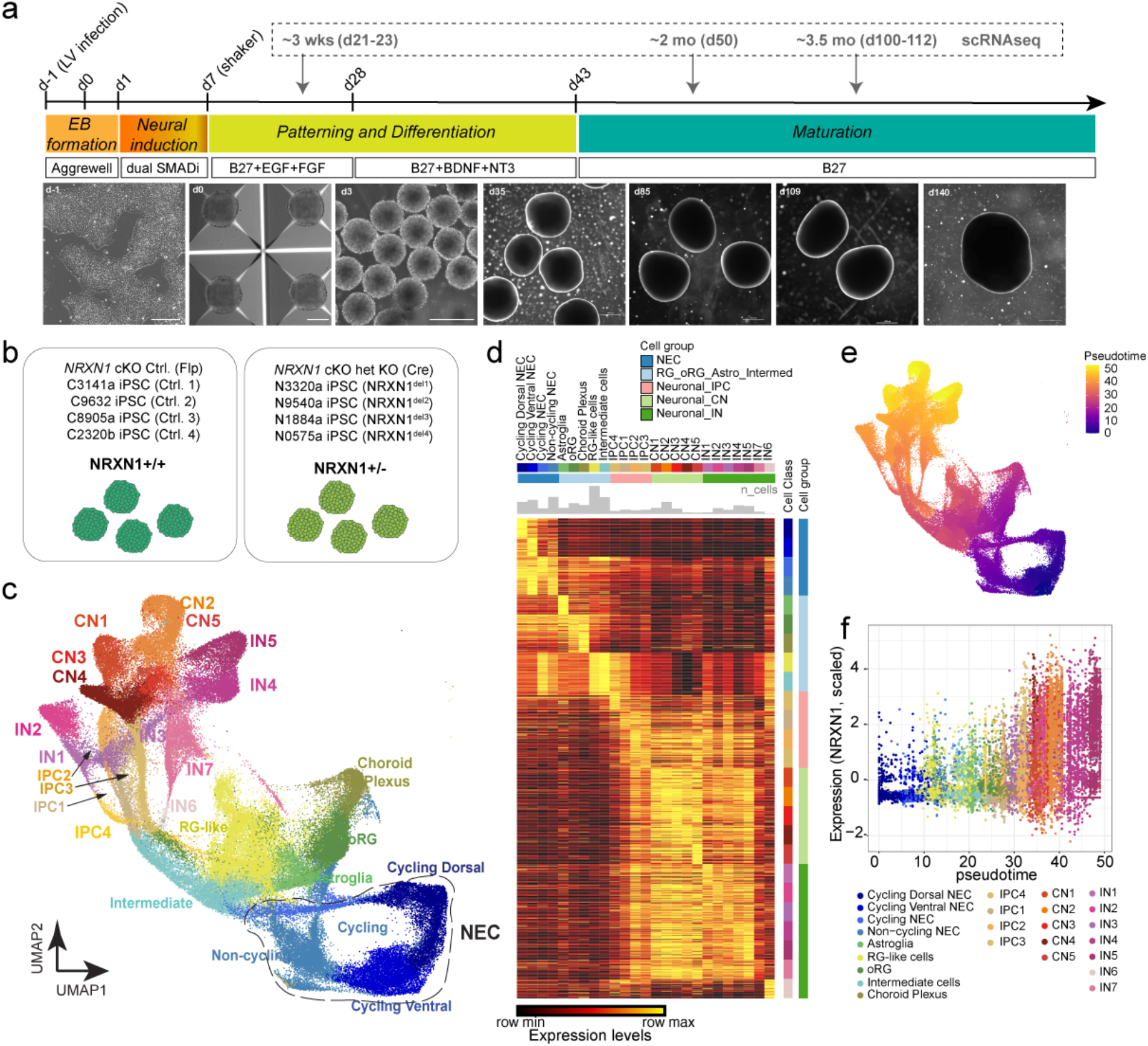
Generation of forebrain organoids from genetically engineered *NRXN1* cKO hESCs and donor derived iPSCs from healthy controls and SCZ patients. (a) Schematic of brain organoid generation protocol and the corresponding representative brightfield images over development. Scale bars – 250 μm for d-1, 0, 3; 100 μm for d35+. (b) Schematic showing genotypes used for scRNAseq and other experiments in the study. (c) Uniform Manifold Approximation and Projection (UMAP) showing distributions of cell classes of the integrated single-cell data. Abbreviations: neural precursor cells (NECs); outer radial glial cells (oRG); radial glial-like cells (RG-like); intermediate precursor cells (IPC); cortical excitatory neurons (CN); and cortical GABAergic inhibitory neurons (IN). (d) Heatmap of gene signatures across cell classes and cell groups. Student t-tests were used to compute differentially expressed genes (DEGs) of each cell class in the integrated scRNA-seq data. Top 50 DEGs for each cell type are shown in a diagonal manner while normalized expression levels are scaled for each row. (e) UMAP showing distribution of pseudotime values of integrated single-cell data inferred by Monocle3. (f) NRXN1 mRNA expression (Seurat scaled expression values) across pseudotime in control donor-derived brain organoids. A higher pseudotime value indicates greater maturity as indicated by the number of various neuronal cell classes.

Having established a reliable protocol, we then subjected a cohort of previously characterized hPSC lines^30, 31^ (Table 1) to brain organoid differentiation and collection for scRNAseq (Fig. 1b). We generated brain organoids from the *NRXN1* cKO hESC line, which produced a single replicate of control (Flp) and deletion (Cre) at day 23, two replicates of control and deletion samples at day 50, and two replicates of control and deletion samples at day d100/112. In addition, two sets of SCZ patient and control donor iPSC pairs (2 SCZ-*NRXN1*^del^ lines and 2 control lines) were sequenced at days 22 and 50 and four sets of SCZ patient and control donor iPSCs (4 SCZ-*NRXN1*^del^ lines and 4 control lines) were sequenced at day 100/101 time point. For simplicity, we designate these time points as ∼3 wk (days 21-23), ∼2 mo (day 50), and ∼3.5 mo (days 100-112), and hereafter refer to *NRXN1* cKO samples as ‘engineered’ and SCZ-*NRXN1*^del^ lines and controls as ‘donor.’ After rigorous quality control data processing steps (Fig. S3-4), we generated a total of 156,966 high-quality single-cell transcriptomes from 26 brain organoid samples for downstream analysis (Fig. S5; n=10 engineered and n=16 donor).

**Table 1.** hPSC lines used for the study.

| Line ID | Name used in this study | Sex | Genotype | Diagnosis | Single-cell RNA sequencing | Calcium Imaging | IHC |
| --- | --- | --- | --- | --- | --- | --- | --- |
| *NRXN1* cKO hESCs | Flp (Ctrl) /Cre (Het) | M | *NRXN1*$\alpha\beta$ deletion | None | yes | yes | yes |
| C3141a iPSC | Ctrl.<sup>1</sup> | M | Control | None | yes | no | no |
| C9632 iPSC | Ctrl.<sup>2</sup> | F | Control | None | yes | yes | yes |
| C8905a iPSC | Ctrl.<sup>3</sup> | M | Control | None | yes | yes | yes |
| C2320b iPSC | Ctrl.<sup>4</sup> | M | Control | None | yes | no | no |
| N3320a iPSC | *NRXN1*<sup>del1</sup> | M | *NRXN1*α deletion | SCZ | yes | no | no |
| N9540a iPSC | *NRXN1*<sup>del2</sup> | F | *NRXN1*$\alpha\beta$ deletion | SCZ | yes | yes | yes |
| N1884a iPSC | *NRXN1*<sup>del3</sup> | M | *NRXN1*α deletion | SCZ | yes | yes | yes |
| N0575a iPSC | *NRXN1*<sup>del4</sup> | F | *NRXN1*α deletion | SCZ | yes | no | no |

## Single cell transcriptomic atlas from developing organoids informs relevant cell types with prominent NRXN1 expression

After data processing, normalization, and clustering, a total of 29 cell clusters were further annotated by marker expression (Fig. S6b, S7), which consisted of both cycling and non-cycling neural progenitors (NECs), radial glia-like cells (RG-like cells), outer radial glial cells (oRGs), astroglia, intermediate cells, and intermediate progenitor cells (IPCs) that give rise to distinct subpopulations of glutamatergic excitatory neurons (CNs) and GABAergic inhibitory neurons (INs; Fig. 1c, d; Fig. S6, S7). We further validated our cell annotations by comparing gene expression signatures of our scRNAseq cell clusters to published brain organoid datasets as reference (Fig. S6a)^20, 33–37^. Remarkably, we saw a high correlation between published cell clusters and ours, suggesting that the specific cell clusters in our scRNAseq dataset share similar gene expression patterns with other brain organoids that were generated with slightly different protocols, thus showing overall reproducibility of 3D culturing across protocols. Here, we also provide an interactive visualization of 3D UMAP for further exploration (see Supplementary .html files).

Using this scRNAseq dataset, we first analyzed the cell-type-specific expression of *NRXN1* gene in the control donor samples across brain organoid development, which would allow identification of specific cell types enriched for *NRXN1* function. We quantified normalized *NRXN1* expression levels and found that CNs and INs reproducibly showed the highest expression of *NRXN1* across the three time points (Fig. S8a). Though not to the same degree as these cell types, IPCs, oRGs, and astroglia did express *NRXN1* at low levels to start (2 mo) and progressively increased in expression over time (3.5 mo). Lastly, NEC subtypes showed the lowest abundance of *NRXN1* (Fig. S8a). In addition, we used developmental trajectory analysis (monocle3^38^) to quantify *NRXN1* expressing cell types across pseudotime (Fig. 1e, f). As expected, using data from the three time points, we saw that the brain organoids underwent a progressive developmental trajectory that mirrored the corresponding maturity across pseudotime: early stage of trajectory corresponding to proliferating cells while later stage of trajectory corresponding to differentiated and mature neuronal subtypes (Fig. 1e, f). Similar to fixed time point analysis, cells with highest *NRXN1* expression included most mature CN and IN subtypes followed by IPC subtypes, oRGs, and astroglia. Based on this result, we concluded that the function of human *NRXN1* gene can be most reliably studied in differentiated neurons and astroglia, as well as in cortical progenitors, such as IPCs and oRGs, in the brain organoid model. While this result highlights the important cell types for *NRXN1* function in organoid models, we were curious about *NRXN1* expression patterns in human primary tissue and similarities with brain organoids. To this end, we performed analysis by leveraging a published human fetal brain scRNAseq dataset, which reported single cell transcriptomes from second trimester microdissected tissues, representing a time period associated with the peak of neurogenesis and early gliogenesis (14 GW to 25 GW)^36^. Comparable to brain organoids, at the youngest fetal age (14 GW), glutamatergic neurons showed the highest normalized *NRXN1* expression, and at 16 GW, both glutamatergic and GABAergic neurons consisted of the highest *NRXN1*-expressing cells (Fig. S8b). These neuronal cells showed comparable expression patterns as progenitor and forebrain radial cells at older time points (20 and 25 GWs). Due to the low number of astrocytes represented in the dataset, astrocytes were not quantified here. In contrast, progenitor cells and radial glial cells initially showed a relatively low number of *NRXN1* expressing cells at 14-16 GW, which reached to similar levels to glutamatergic neurons and GABAergic neurons at 20-25 GW (Fig. S8b). Thus, based on both organoid and human tissue data, major cell types, including glutamatergic neurons and GABAergic neurons in addition to forebrain progenitor cells and astroglia, may be most affected by *NRXN1* haploinsufficiency in the developing forebrain.

## NRXN1 engineered deletions affect mature time point and alter gene expression programs in ubiquitin proteasome system, alternative splicing, and synaptic signaling

To study the impact of developmental-timing- and cell-type-specific effects of isogenic *NRXN1* deletions in brain organoid development, we analyzed the engineered samples throughout the three time points by 1) constructing single-cell developmental trajectories across pseudotime and 2) analyzing cell-type-specific differentially expressed gene (DEG) networks. By calculating the densities of cells across pseudotime values, we drew density plots for cells with (Cre) or without engineered *NRXN1* deletions (Flp) across three time points, representing the dynamic cell abundance changes and cellular transitions throughout their developmental trajectory (Fig. 2a,b). Interestingly, we found that engineered deletions followed similar developmental trajectories as controls at 3 wk and 2 mo until reaching 3.5 mo, which showed a subtle but noticeable difference at two peaks between the control and deletion (Fig. 2b). Engineered deletion organoids displayed deviations in developmental trajectories impacting RG-like cells, oRGs, and astroglia (first peak) and IPCs and CNs/INs (second peak), suggesting that the timing of cellular differentiation and maturation in brain organoids may be affected. Importantly, at 3 to 4 month time period, brain organoids reach the peak of neuronal diversity and amplification and the beginnings of astrogenesis^19, 33, 39^, indicating that engineered *NRXN1* deletions may impact gene expression programs that regulate active neurogenesis, gliogenesis, and synapse development.

**Figure 2.**
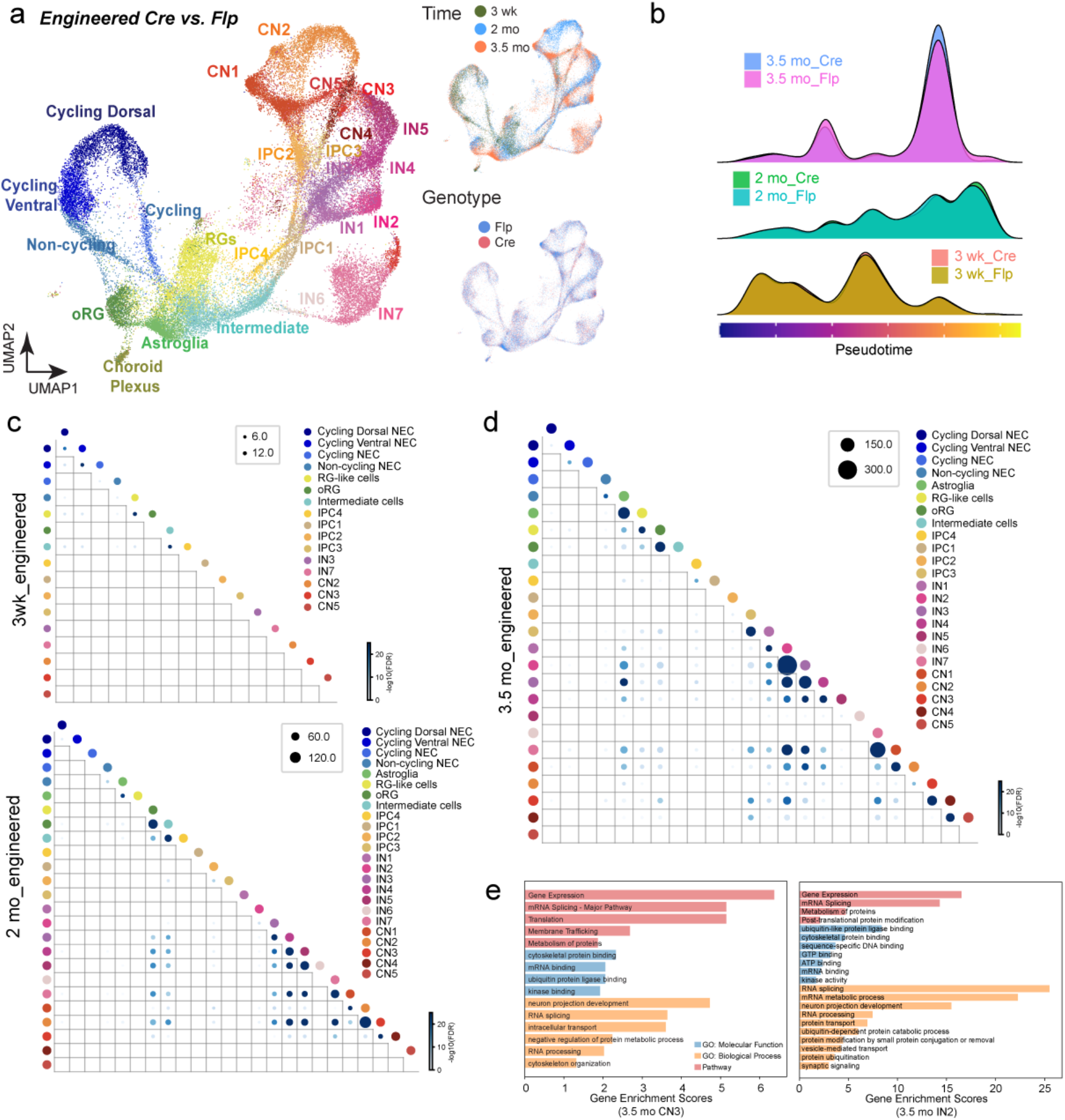
Perturbation effects of *NRXN1* engineered deletions across development. (a) UMAPs showing distributions of cell classes (left), time points (top right), and genotypes (bottom right) of *NRXN1* cKO engineered brain organoids. A total of 10 samples were processed for deletion (Cre) control (Flp) – n=1 for 3 weeks (3 wk) Cre and Flp; n=2 for 2 months (2 mo Cre and Flp; n=2 for 3.5 months (3.5 mo) Cre and Flp. (b) Ridge plots showing the density of cell abundance across the dimension of pseudotime for 3 wk, 2 mo, and 3.5 mo engineered brain organoids. (c & d) Dot plots showing significant differential gene expression in deletion vs. control across time points and cell types. The size and color of each dot show the number and significance of overlapping DEGs for each comparison of two cell classes. The significance was measured by −𝑙𝑜𝑔_10_(𝐹𝐷𝑅 𝑎𝑑𝑗𝑢𝑠𝑡𝑒𝑑 𝑝 𝑣𝑎𝑙𝑢𝑒𝑠) of hypergeometric tests (see Methods). (e) Representative enriched gene sets were shown for GSEA results of DEGs of multiple cell classes using ToppGene. Different categories of Gene Ontology are shown in different colors (blue: molecular function, orange: biological process, and red: pathway). Enrichment scores were defined as −𝑙𝑜𝑔_10_(𝐹𝐷𝑅 𝑎𝑑𝑗𝑢𝑠𝑡𝑒𝑑 𝑝 𝑣𝑎𝑙𝑢𝑒𝑠) to represent the associations between DEG sets and Gene Ontology gene sets.

To explore DEG networks in engineered deletions vs. controls, we performed DEG analysis (FDR-adj. p< 0.05) in each cell type associated with each time point. Total DEG number increased from 2 mo to 3.5 mo with the greatest perturbation at 3.5 mo (Fig. 2c-e, Table S2). In 3.5 mo engineered organoids, DEG patterns were most pronounced in astroglia, RG-like cells, oRGs, IPC3, CNs, and INs (Fig. 2c-e, Table S2), which correlated with the trajectory analysis (Fig. 2b). To better understand the contribution of specific gene modules associated with cell-type-specific DEGs across each developmental time point, we explored whether there exists any overlap between DEGs in different cell types using a hypergeometric test. Interestingly, DEG overlapping patterns were present within CNs and INs at 2 mo and 3.5 mo, showing similar DEGs impacted within neuronal subtypes (Fig. 2d, Table S3).

Using 3.5 mo cell-type-specific DEGs, we performed gene set enrichment analysis (GSEA, ToppGene^40^) to examine whether specific molecular functions, biological processes and/or biochemical pathways were significantly enriched. Collectively, DEG sets from astroglia, oRGs, IPC3, CNs, and INs were specifically enriched for RNA splicing/processing and Ubiquitin (Ub)-mediated processes (Fig. 2e, S9, Table S4). Notably, DEGs representing both glutamatergic excitatory neurons (CN3/4) and GABAergic neurons (IN2/7) were enriched for synaptic genes as expected, as well as components of the Ub-mediated proteolysis and RNA splicing (Fig. 2e, S9, Table S4). Intrigued by the specificity of the DEG enrichment in the components of mRNA splicing and ubiquitin-proteasome system (UPS), we further queried for these gene signatures across time points and cell types. Splicing- and UPS-specific DEGs are detected early as 2 mo, and by 3 mo, the impact of these DEGs became more enhanced (Fig. S10, 11), suggesting that perturbation effects become amplified as cells mature. Altogether, GSEA suggests that regulators of RNA splicing and UPS are consistently perturbed from oRGs to differentiated neuronal subtypes in the engineered deletions.

## SCZ-NRXN1 deletions induce early developmental perturbation effects which are maintained throughout maturation with consistent DEG patterns as engineered deletions

Using similar analysis methods, we probed how SCZ-*NRXN1* deletions affect single-cell developmental trajectories and DEG networks. To our surprise, we found a significant and consistent deviation of pseudotime developmental trajectories from 3 wk to 3.5 mo in the donor deletions vs. controls (Fig. 3a,b). Most notably, donor deletions induced abnormal developmental patterns of NECs in addition to other developing cell types, an effect that was not observed in the engineered deletion organoids. Mirroring this finding, we observed significant DEG occurrences in NECs from 3 wk time point, which was maintained up to 3.5 mo (Fig. 3c,d, Table S3, S5). At 3.5 mo, there was a great degree of DEG overlaps across cell types, indicating possible overlap of common biological effects. In summary, the DEG significance, overlapping DEG patterns, and abnormal developmental trajectories were all enhanced in the donor deletions compared to the engineered deletions.

**Figure 3.**
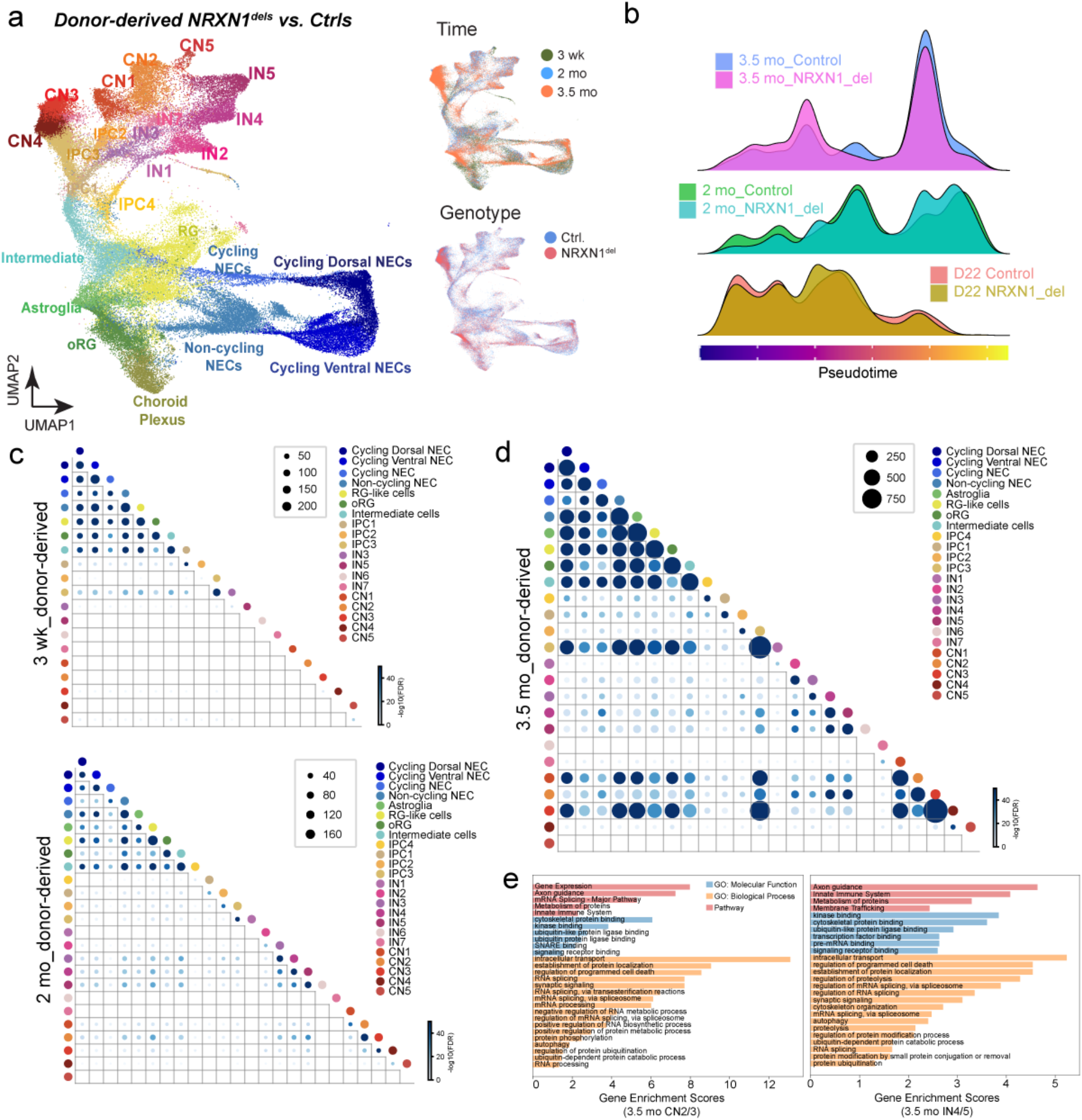
Perturbation effects of SCZ associated *NRXN1* deletions across development. (a) UMAPs showing distributions of cell classes (left), time points (top right), and genotypes (bottom right) of SCZ-*NRXN1*^del^ donor derived brain organoids. A total of 16 samples were processed – 2 SCZ-*NRXN1*^del^ donors and 2 Ctrl donors for 3 weeks (3 wk); 2 SCZ-*NRXN1*^del^ donors and 2 Ctrl donors for 2 months (2 mo); 4 SCZ-*NRXN1*^del^ donors and 4 Ctrl donors for 3.5 months (3.5 mo). (b) Ridge plots showing the density of cell abundance across the dimension of pseudotime for 3 wk, 2 mo, and 3.5 mo engineered brain organoids. (c & d) Dot plots showing significant differential gene expression in deletion vs. control across time points and cell types. The size and color of each dot show the number and significance of overlapping DEGs for each comparison of two cell classes. The significance was measured by −𝑙𝑜𝑔_10_(𝐹𝐷𝑅 𝑎𝑑𝑗𝑢𝑠𝑡𝑒𝑑 𝑝 𝑣𝑎𝑙𝑢𝑒𝑠) of hypergeometric tests (see Methods). (e) Representative enriched gene sets were shown for GSEA results of DEGs of multiple cell classes using ToppGene. Different categories of Gene Ontology are shown in different colors (blue: molecular function, orange: biological process, and red: pathway). Enrichment scores were defined as −𝑙𝑜𝑔_10_(𝐹𝐷𝑅 𝑎𝑑𝑗𝑢𝑠𝑡𝑒𝑑 𝑝 𝑣𝑎𝑙𝑢𝑒𝑠) to represent the associations between DEG sets and Gene Ontology gene sets.

We then investigated the functional significance of DEGs in donor deletions and whether they represent similar or different gene set enrichment patterns compared to engineered deletions. Therefore, we performed GSEA on overlapping DEGs found in similar cell types organized into separate cell groups – all NECs, RG-like cells/intermediate cells/IPC3, astroglia, oRGs, CN2/3, and IN4/5. Intriguingly, while each cell group showed specific DEG enrichment of biological processes relevant to its cell type functions, i.e. cell cycle-related genes in dividing cells and cell signaling genes in neurons, we again identified two distinct biological processes that were repeatedly observed across cell groups – RNA splicing and UPS (Fig. 3e, S12, Table S6). As mentioned above, these two pathways were also found disrupted in engineered deletions. Specifically in neuronal subtypes (CN2/3 and IN4/5), a strong enrichment of synaptic signaling genes was present (Fig. 3e, S12, Table S6). We next tested for the presence of RNA splicing- and UPS-specific DEGs in all cell types across timepoints and found that donor deletions showed a significant enrichment of these DEGs throughout development with most significant patterns reached at 3.5 mo (Fig. S10, S11). Collectively, these results highlight that, in the developing brain organoids, gene networks involving UPS and RNA splicing are commonly disrupted across isogenic and SCZ-*NRXN1* deletion backgrounds but manifest in different temporal and cell-type-specific manners.

## Alternative splicing dysregulation in glutamatergic excitatory neurons with NRXN1 deletions

To gain better insight into how RNA splicing could be misregulated in our brain organoid models, we examined the expression patterns of established neuronal splicing regulators including PTBPs, MBNLs, RBFOXs, NOVAs, SSRMs, and KHDRBSs in both the donor and engineered deletion contexts at 3.5 mo (Table S7). Interestingly, we found that the vast majority were downregulated, suggesting that these splicing regulators may not function properly to promote global splicing patterns of mature neuronal programs^41–45^. Based on the predicted functions of these RNA binding proteins in regulating alternative splicing during neural development and their associated downregulated gene signatures, we reasoned that *NRXN1* deletions could render an overall immature splicing program and that global alternative splicing could potentially be misregulated. Moreover, since many of these neuronal splicing regulators control alternative splicing of neurexins^46^, we questioned whether alternative splicing of *NRXN1* gene itself was changed as previously reported in postmortem tissue samples from SCZ patients^47^.

To test whether *NRXN1* deletion induces changes in alternative splicing programs in neurons, we explored a previously published bulk RNA sequencing dataset generated from mature glutamatergic excitatory induced neurons (iNs) from the same set of donor derived iPSC lines and an engineered *NRXN1* cKO iPSC line^31^. We first confirmed that neuronal splicing regulators are differentially expressed with an overall downregulation pattern in iNs with *NRXN1* deletion (Fig. 4a). Using RSEM^48^, we detected a total of 184 differentially expressed isoforms (DEIs) enriched in engineered iNs with *NRXN1* deletions (Cre/Flp) and 213 DEIs in SCZ donor iNs with *NRXN1* deletions (*NRXN1*^dels^/Ctrls) across three pairs of healthy control and SCZ-*NRXN1* deletion carriers (Fig. 4b, Table S8). These DEIs mapped to 165 and 153 unique gene IDs for engineered iNs and donor iNs, respectively. Among the differentially spliced genes, ∼10% consisted of synaptic genes identified by SYNGO enrichment analysis^49^, implicating a major functional impact on synaptic signaling. Interestingly, while 5 genes (*AHI1, CHD4, DHX36, NRXN1, STAMBP*) overlapped between engineered and donor iNs, the majority of genes were unique to each iN type (Fig. 4b), showing differential abundance of DEIs in the engineered vs. donor genetic background. Interestingly, synaptically localized *CHD4* and *NRXN1* showed similar differential expression patterns of isoforms in both deletion contexts (Fig. 4c,d).

**Figure 4.**
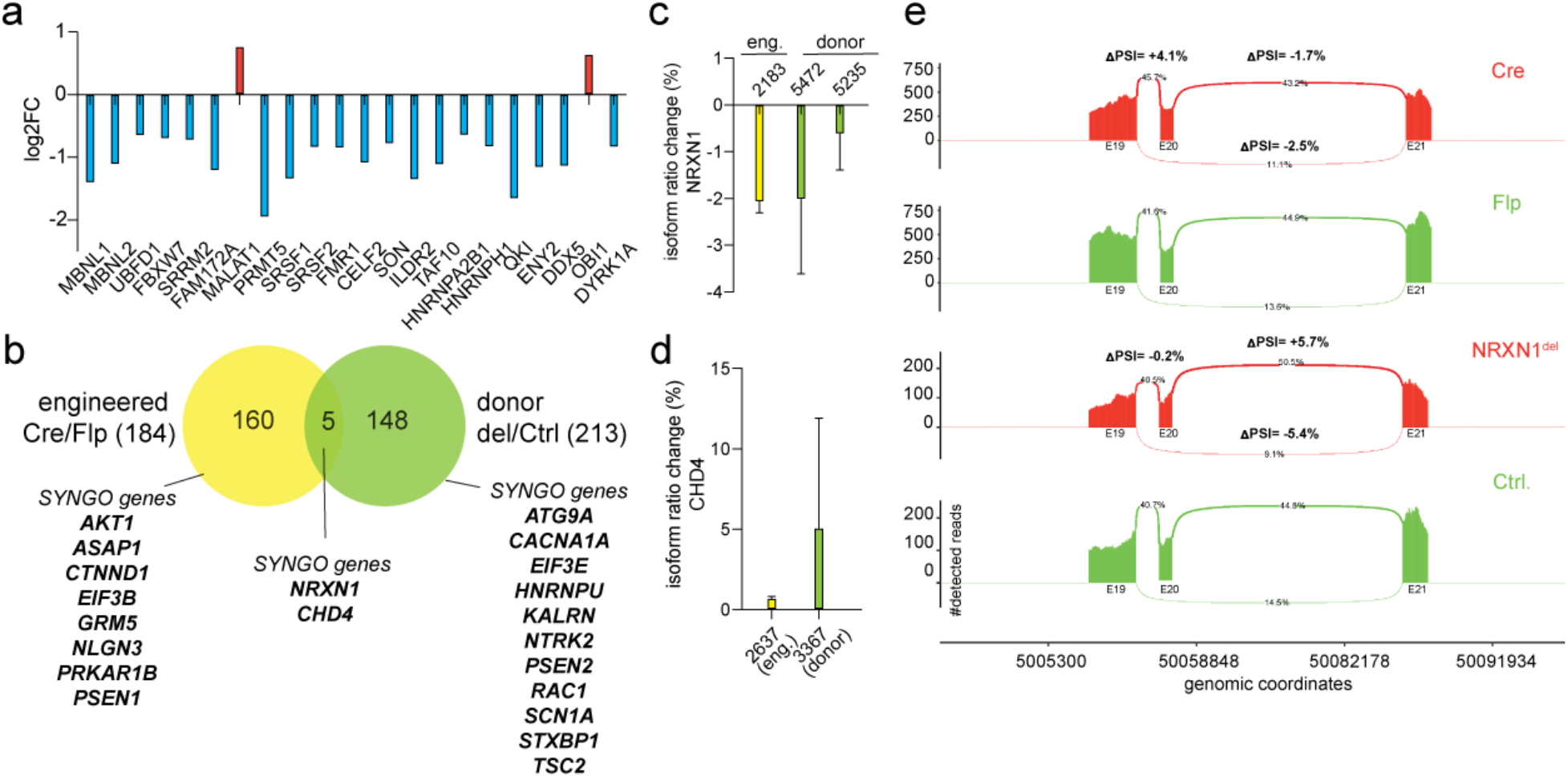
Dysregulated alternative splicing in *NRXN1* mutant glutamatergic human neurons. (a) Differential gene expression (adjusted p values < 0.05) of neuronal splicing regulators in SCZ-*NRXN1*^del1^ compared to Ctrl.^1^ iN cells (n=3 replicates). Log_2_ fold changes are plotted on the y-axis with example DEGs on the x-axis. Blue bars represent downregulation and red bars represent upregulation. (b) Venn diagram representing differential isoform abundance derived from RSEM analysis across iN cells from engineered *NRXN1* deletions (n=3 replicates from *NRXN1* cKO iPSC) and donor-derived SCZ-*NRXN1* deletions (3 healthy controls and 3 SCZ-*NRXN1* deletion carriers; n=3 replicates per line, 18 replicates total). Number of differentially expressed isoforms (184 for engineered iNs and 213 donor derived iNs) mapped to 160 and 148 unique gene IDs respectively. Among the unique genes, example SYNGO-mapped genes are highlighted. (c-d) Percentage of isoform ratio change (differential transcript usage, see methods) for *NRXN1* and CHD4 in *NRXN1* deletions from engineered (eng.; yellow) and donor (green) backgrounds. (e) Detection and visualization of local differential splicing using Majiq and ggSashimi. Change in percent spliced in index (dPSI) is calculated for differential exon usage in the *NRXN1* gene (exons 19-21) in the SCZ-*NRXN1* deletion vs. *NRXN1* cKO engineered deletion iNs. Number of detected reads is plotted on the y-axis with location of genomic coordinates on the x-axis.

In addition to differential isoform expression, we analyzed differential local splicing in iNs using Majiq^50^. In this analysis, one can integrate both *de novo* splicing change detection as well as known splicing events to quantify the change in PSI (percent spliced in index), which reports how often specific sequences are spliced into transcripts. Out of 33 specific local splicing variations (LSVs) present in the *NRXN1* gene in iNs, we found 4 LSVs which were similarly changed in both donor and engineered iNs, 15 LSVs that were different, and 14 LSVs that were not significantly changed in any genotype (Table S9). More specifically, we found a biologically significant LSV in exon 20, where inclusion of this exon was increased in the deletions compared to controls in both isogenic and SCZ-*NRXN1* deletion backgrounds (Fig. 4e). Differential splicing of this exon, which contains splice site #4, could have functional implications in ligand binding and synaptic properties^5, 7^. Altogether, splicing analysis of mature glutamatergic iNs show that *NRXN1* deletions result in changes in spliced variant landscape of synaptic genes, including *NRXN1* itself, which could impact synaptic function by differential isoform representation.

## Impaired neuronal network function in brain organoids carrying NRXN1 deletions

To test whether the observed developmental abnormalities and gene expression programs translate to functional and sustained differences in neuronal connectivity, we performed live Ca^2+^ imaging in 4 mo brain organoids from SCZ-*NRXN1* deletion vs. control (*NRXN1*^del2^ vs. Ctrl^2^). Two weeks prior to imaging, brains organoids were infected with AAVs expressing a genetically encoded Ca^2+^ indicator, somaGCaMP6fs construct^51^. Under basal conditions (without any stimulation), we measured the frequency and amplitude of spontaneous Ca^2+^ transients, which are indicative of spontaneous neuronal network activities. In addition, we quantified the frequency of synchronous firing, which indicates how often neurons fire together, thereby producing synchronized bursts of activities. Compared to controls, *NRXN1*^del2^ brain organoids showed a significant decrease in the frequency of spontaneous Ca^2+^ transients without a change in the amplitude of the responses, as measured by dF/F_0_ intensity (Fig. 5a-c). In addition, there was an overall decrease in the synchronous firing rate in these brain organoids, demonstrating a significant decrease in the neuronal network bursts (Fig. 5a-c).

**Figure 5.**
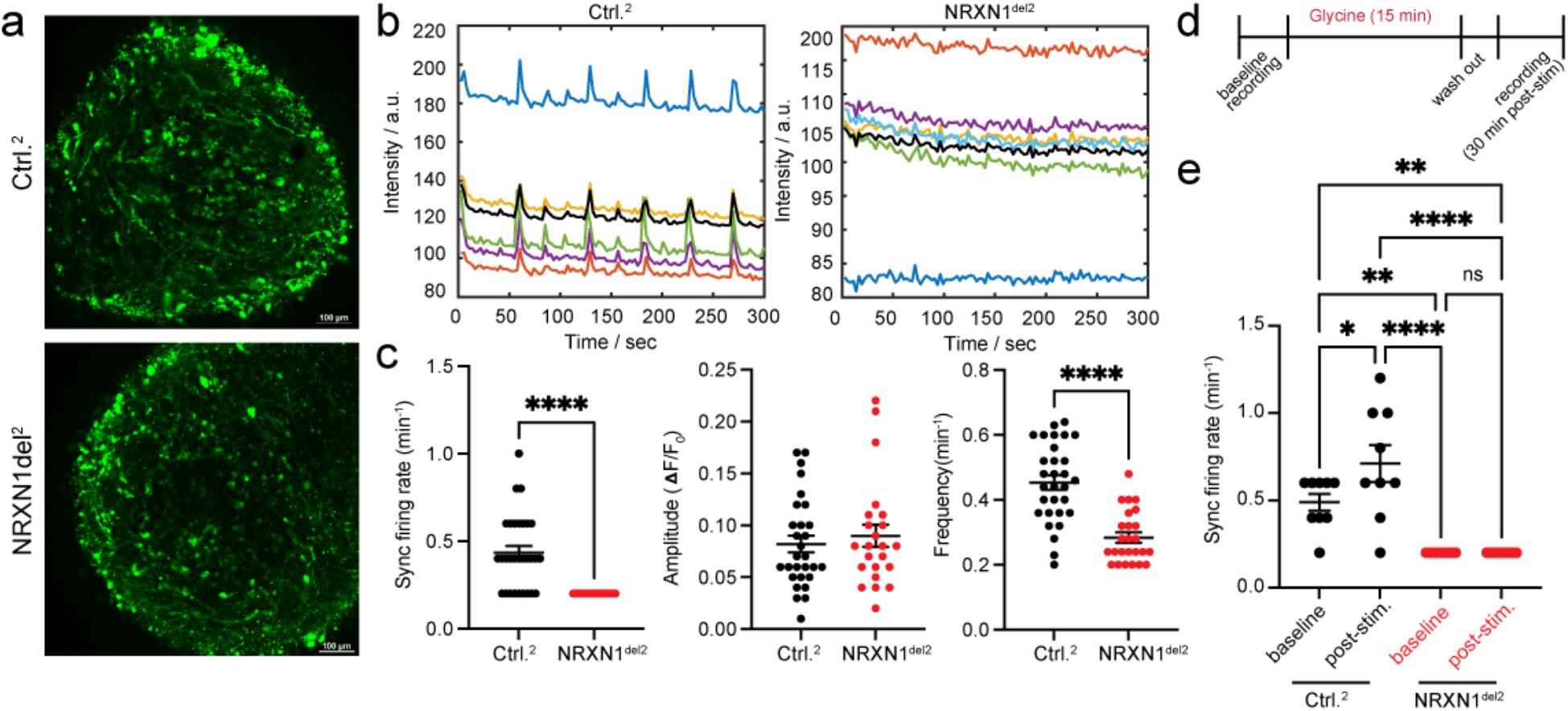
Impaired neuronal network activities in brain organoids carrying *NRXN1* deletion. (a) Representative fluorescent images of intact donor derived organoids (Ctrl.^2^ and *NRXN1*^del2^) expressing soma-GCaMP6f2 at ∼120 days prior to live Ca^2+^ imaging. (b) Colored raw intensity traces are shown in the boxed graphs with averaged intensities plotted in bolded black. (c) Averaged data for synchronous firing rates (number of detected synchronous spikes/minute) representative of network activity, as well as amplitudes (dF/F_0_) and frequencies (total number of detected peaks/minute) of spontaneous spike activity, are shown in scatter plots. Each data point represents averaged data from a single field of view (FOV) consisting of 4-5 ROIs per FOV. At least 4-6 FOVs were taken from each organoid and 5 organoids per genotype were used for experiments. (d) Experimental flow of live Ca^2+^ imaging with glycine stimulation. (e) Synchronous firing rate is plotted in genotypes with or without stimulation. Each data point represents averaged data from a single field of view (FOV) consisting of 4-5 ROIs per FOV. At least 4-6 FOVs were taken from each organoid and 2 organoids per genotype were used for experiments. Error bars represent S.E.M. Statistical significance is represented by asterisks: *p < 0.05, **p < 0.01, ***p < 0.001, and ****p < 0.0001. Student’s t-test was performed for data shown for (c) and one-way ANOVA with multiple comparisons (Tukey’s test) was used for data shown on (e).

Next, we explored the possibility that N-methyl-D-aspartate receptor (NMDAR) function might be abrogated in these organoids, thus contributing to the overall neuronal network dysfunction. Based on DEG analysis of 3.5 mo brain organoids, we identified differential expression in NMDAR subunit 2B (*GRIN2B*) and fatty acid binding protein 7 (*FABP7*), both of which have been linked to SCZ pathogenesis^52–56^. It is well documented that NMDAR hypofunction underlies SCZ pathology^57, 58^, and recent exome sequencing and GWAS studies identified the NMDAR subunit *GRIN2A* as a significant SCZ risk allele^59^. Genetic variants in *FABP7* have been identified in SCZ and ASD patients, and its function has been linked to NMDAR signaling regulation^52–54, 60^. Furthermore, while searching for neuronal-specific DEGs that were consistently perturbed between engineered and donor deletions, we found *GRIN2B* as a commonly perturbed gene in GABAergic neuronal subtypes across genetic backgrounds (engineered IN2 vs. donor IN4/5; Tables S2, S5). These findings suggest that misregulated NMDAR signaling in GABAergic neurons could potentially impact synaptic connectivity and signaling in these brain organoid models. To test this idea, we developed an assay mimicking chemical long term potentiation (cLTP) whereby the functionality of NMDARs can be tested with the addition of glycine, a co-agonist for NMDARs^61^ (Fig. S13a). In control brain organoids, glycine stimulation potentiates neuronal activity as shown by sustained increases in synchronous firing rate measured at 30 min and 60 min post-stimulation (Fig. S13b, d, e). This potentiated activity is mediated by NMDARs, as the addition of selective NMDAR antagonist APV blocks this effect (Fig. S13c-e). Indeed, when stimulated with glycine, brain organoids with SCZ-*NRXN1* deletion failed to potentiate glycine-induced activity compared to control brain organoids (Fig. 5e), suggesting disruption in NMDAR activity.

At more mature time points, (∼4.5 – 5 months), we found that both synchronous neuronal activity as well as spontaneous neuronal spikes was still decreased in the SCZ-*NRXN1* deletions (*NRXN1*del^3^) compared to controls (Fig. S14a-d), suggesting synaptic network dysfunction continues through maturation in these models. Interestingly, *NRXN1* engineered deletions produced a slightly different phenotype in which the frequency of spontaneous Ca^2+^ transients was increased without any changes in the amplitude of the responses as well as the synchronicity of spontaneous firing events (Fig. S14e-h). Recorded Ca^2+^ transients represent true neuronal signals as they were sensitive to synaptic blockers (Fig. S15). These data suggest that although spontaneous neuronal activities are uniformly altered in the brain organoids carrying *NRXN1* deletions, depending on the genetic background, different phenotypic outcomes manifest, reflective of the differences in transcriptomic landscape and developmental origins of such changes in these brain organoids.

## Differential enrichment of disease-associated signatures in SCZ-NRXN1 deletions vs. engineered NRXN1 deletions

To test whether up- and down-regulated DEGs identified from donor-derived and engineered brain organoids were associated with specific neuropsychiatric disease gene signatures, we computed a ‘disease enrichment’ score (-log10 (FDR-adjusted *p* values)) based on a previously established curated list of dysregulated gene sets obtained from SCZ, bipolar disorder (BD), major depression disorder (MDD), and ASD-associated samples^62^. Excitingly, in 3.5 mo brain organoids, DEGs from both donor and engineered deletions were significantly enriched for ASD and SCZ gene signatures (Fig. 6a). Interestingly, there was no significant enrichment of the DEGs in MDD- and BD-related gene sets across the cell types, suggesting that the DEG pool from our brain organoids most closely resembles dysregulated transcriptional signatures related to SCZ and ASD, similar to what has been reported regarding shared genetic signals between SCZ and ASD^63^.

**Figure 6.**
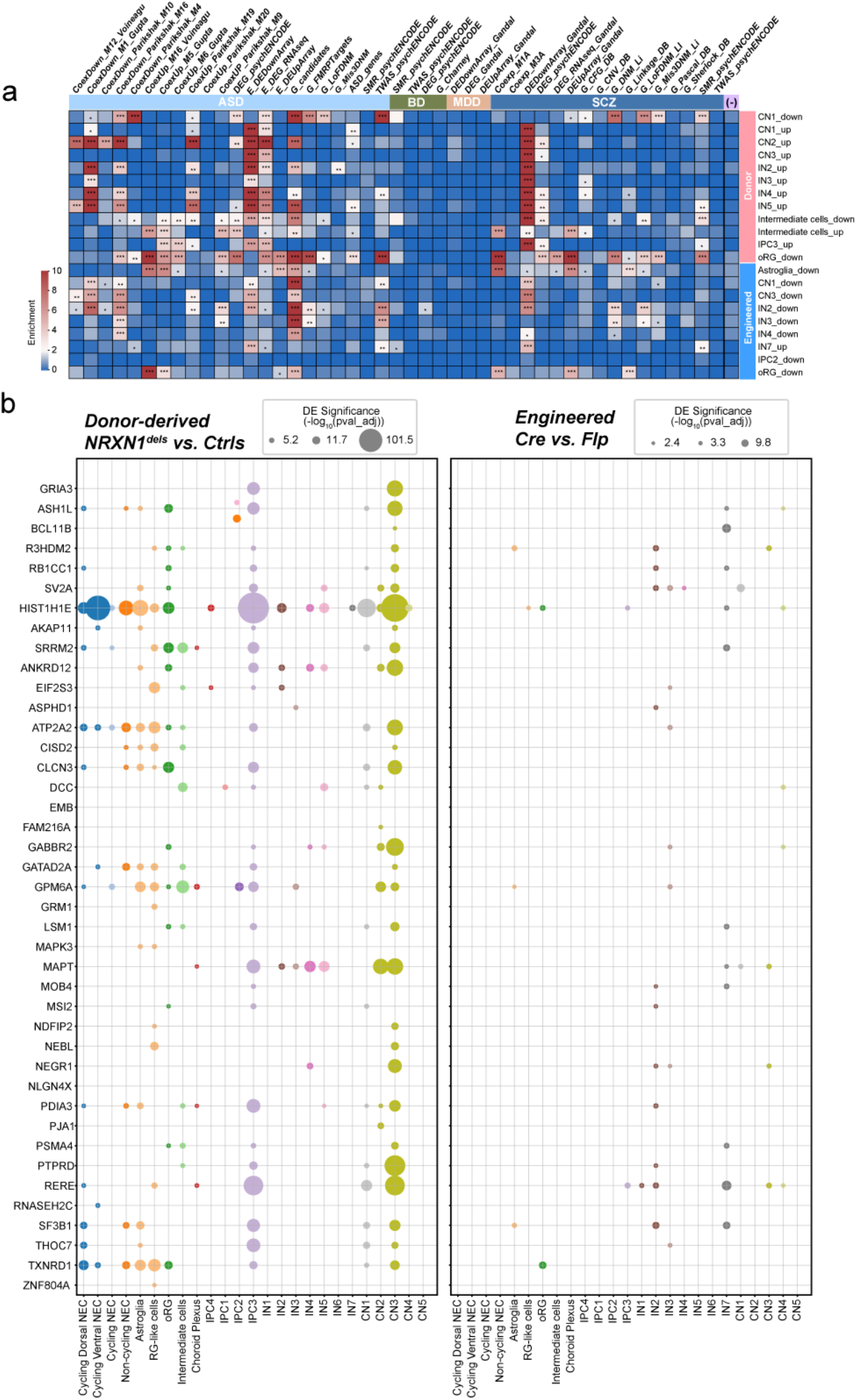
Differential enrichment of disease-associated transcriptomic signatures. (a) Heatmap showing results from a subset of gene enrichment analyses of DEGs performed on cells of both brain organoid types (engineered and donor derived brain organoids) across all time points using neurological disorder dysregulated gene sets in several categories (autism spectrum disorders, ASD; bipolar disorder, BP; mood disorder, MDD; schizophrenia, SCZ) (columns). Data shown here are from 3.5 mo time point only. Body Mass Index (BMI) was used as control (-). Significance scores were defined as −𝑙𝑜𝑔_10_(𝐹𝐷𝑅 𝑎𝑑𝑗𝑢𝑠𝑡𝑒𝑑 𝑝 𝑣𝑎𝑙𝑢𝑒𝑠) to represent the associations between DEG sets and neurological disorders. Scores were trimmed to 0∼10 (see Methods). Significance levels were represented by numbers of asterisks (*: adjusted p values < 0.05; **: adjusted p values < 0.01; ***: adjusted p values < 0.001). (b) Significance of differential expression of prioritized genes obtained from PGC wave 3 and SCHMEA consortium^51, 63^. The size of each dot represents the level of DE significance of each gene in each cell class of 3.5 mo donor brain organoids (left; n=4 for Ctrls and n=4 for SCZ-NRXN1^dels^) and 3.5 mo engineered brain organoids (right; n=2 for Flp and n=2 for Cre).

Next, we independently examined the enrichment score of rare and common variants of SCZ in the 3.5 mo organoid DEG sets by comparing them to a list of risk genes recently reported by SCZ-GWAS PGC wave3 and SCHEMA consortium^59, 63^. Remarkably, several of the SCZ risk genes were represented across cell types in the SCZ-*NRXN1* deletions (Fig. 6b), which highlights SCZ-specific transcriptional signatures present in the patient genetic backgrounds. Engineered *NRXN1* deletion-associated DEGs showed minimal overlap with SCZ-associated risk variants, clearly demonstrating strong association of SCZ-specific loci in the patient disease background compared to the isogenic background (Fig. 6b). A separate survey of DEG overlaps with the GWAS-loci across neuropsychiatric disorders^64–67^ confirmed this finding, where we observed the presence of many more disease-associated DEGs in the donor vs. engineered deletions (Table S10).

## Discussion

Here we provide a systematic analysis of the developmental-timing- and cell-type-dependent perturbations induced by *NRXN1* deletions in the developing human brain organoids using single cell transcriptomics. We initially had two specific goals in mind – 1) to understand the developmental effects of *NRXN1* heterozygous deletions in an isogenic background and uncover which time points and cell types are important for *NRXN1* function, and 2) to utilize SCZ-*NRXN1*^del^ patient iPSC-derived organoids as a model to study the molecular and cellular biology of SCZ. By profiling the single cell transcriptomes of *NRXN1* cKO brain organoids, we found that cellular phenotypes associated with *NRXN1* haploinsufficiency manifest during a developmental window of brain organoids at the peak of neurogenesis and early astrogenesis. Moreover, developmental trajectories and gene expression profiles of maturing glutamatergic and GABAergic neurons are impacted by *NRXN1* deletions.

By comparing engineered and donor-derived organoids side by side, we found both commonalities and differences, reflective of the contribution of genetic background effects. We unbiasedly found shared molecular programs that are perturbed across organoid types including alternative splicing, UPS, and synaptic signaling. Synaptic signaling misregulation was expected since *NRXN1* deletions in 2D glutamatergic iNs decrease probability of neurotransmitter release and synaptic strength^31^. In our study, NMDAR subunit *GRIN2B* was differentially expressed in GABAergic neurons across genetic backgrounds (in engineered IN2 and donor IN4/5). This finding is significant as it confirms previous results showing that mouse *Nrxn1* signals through NMDARs^7, 8^ and SCZ-*NRXN1*^del^ glutamatergic iNs carry upregulated levels of the endogenous NMDAR antagonist *KYAT3*^31^. Moreover, genetic variants in the NMDAR subunits, *GRIN2B* and *GRIN2A*, are both observed in SCZ populations^59, 68, 69^. Lastly, NMDAR hypofunction in SCZ has been a longstanding hypothesis supported by multiple post-mortem studies and brain imaging studies from SCZ patients as well as mouse models of NMDAR blockade through ketamine and phencyclidine^57, 58^. In agreement with transcriptomic data, we showed that SCZ-*NRXN1^del^* organoids fail to potentiate glycine-induced augmentation in neuronal activity (i.e. cLTP), which validates NMDAR dysfunction.

Alternative splicing is highly regulated in the brain^70, 71^ and is influenced by neuronally enriched splicing factors like NOVAs, PTBPs, RBFOXs, and MBLNs. The general notion is that RBFOXs and MBNLs promote mature splicing patterns whereas MBNLs antagonize this process, altogether tightly regulating alternative splice usage during cortical development and plasticity^41–45, 72^. Interrogation of DEGs in our brain organoid dataset showed a clear misregulation of these neuronal-specific splicing regulators at the transcriptional level, suggestive of an immature splicing program in these models. Indeed, by analyzing splicing variation in 2D glutamatergic iNs differentiated from the same set of iPSC lines revealed changes in alternative splicing in the abundance of isoform representation and local splicing variation in both donor and engineered deletions. It has been shown that global splicing changes and alternative transcript usage are overrepresented in SCZ brains, more so than in ASDs and BD^47^. Differential splicing of various genes has been observed in the brain samples of SCZ patients compared to controls, including *DRD2, NRG1, ERBB4, GRM3,* and *GRIN1*^47, 73–76^. More recently, differential splicing effects of *NRXN1* has been suggested in SCZ iPSC-derived neurons^77^ and in postmortem brains of SCZ and BD patients^14, 47^, further highlighting the importance of splicing regulation in SCZ as a potential molecular mechanism.

In addition to splicing, DEGs responsible for UPS regulation have been identified in our study. These genes encode for proteins that are E2 conjugating enzymes (*UBE2M, UBE2V2*), deubiquitinase (*UCHL1*), Ub processing-(*UBB*) or autophagy-related factors *(GABARPL2*) as well as proteins, which are themselves or direct binding partners to E3 Ub ligases (*NDFIP1, SKP1, RNF, STUB1*). Though it is not clear from the list of DEGs whether or not these molecules are actively participating in the protein quality control or in the regulation of protein components involved in signal transduction, it has been hypothesized that alterations in proteostasis and Ub-mediated regulation of synaptic signaling contribute to SCZ pathogenesis^78–82^. Additionally, protein truncating variants of Ub ligases (*CUL1* and *HERC1*) were recently found to be associated with SCZ at exome-wide scale^63^, further highlighting the importance of this molecular pathway in SCZ. Possibly more UPS genes are to be discovered for both rare and common variants in the future. Though it remains to be determined whether these changes are indeed causal or merely reporting a consequence of the disease, altered UPS does exist and this, in turn, could affect protein homeostasis in the brains of SCZ patients. Importantly, proteasome function at synapses is tightly regulated by NMDAR activity, as NMDAR activation regulates 26S proteosome assembly and catalytic activity^83, 84^ and stability of proteasomes in the post-synaptic density^85^. Moreover, E3 Ub-ligases and deubiquitinases act in concert to regulate the ubiquitination, internalization and localization of NMDARs, AMPARs, and mGluRs, and therefore, actively participate in Hebbian and homeostatic plasticity^86–88^. Further investigations on the interplay between NMDAR signaling and UPS regulation during synaptic development would enhance our understanding of how these distinct biological pathways converge in the context of SCZ pathogenesis.

Interestingly, there were major differences between donor vs. engineered organoids that we observed. First, unlike the donor derived organoids, engineered organoids did not exhibit changes in the developmental trajectory or gene expression in the NEC subtypes. Moreover, the magnitude of gene expression changes in various cell types in the engineered organoids was minimal compared to donor derived organoids. These findings indicate that brain organoids derived from patient genetic background induce a greater degree of transcriptional perturbations and uncover NECs as a vulnerable cell type during cortical development. Neuronal subtypes, astroglial cells, and oRGs are commonly affected to varying degrees in both patient and engineered genetic backgrounds carrying *NRXN1* deletions. The specificity of NEC phenotypes in the SCZ-*NRXN1*^del^ genetic background is further supported by previous studies reporting alterations in the morphology, differentiation potential, and gene expression profiles from SCZ iPSC derived NECs and brain organoid models, all of which are in support of the neurodevelopmental hypothesis of SCZ^89–91^. Second, donor derived organoids showed a greater accumulation of disease-specific signatures related to SCZ. This finding makes sense, since donor derived organoids carry SCZ-relevant genetic background. Due to this effect, differences in the magnitude of gene expression changes and the directionality of those changes were observed in these organoid types. This is also apparent in the differences in the specific neuronal firing patterns observed in the isogenic engineered vs. donor derived organoids.

There are two main limitations to this study. First, despite obtaining high-quality dataset of ∼157,000 single cells, the overall study is underpowered due to the small sample size of patient/control cohort with limited genetic backgrounds being represented. Larger sample size and multiple technical replicates would allow for a more granular analysis of mutational effects across brain organoid development. Second, while we focused on early developmental time points leading up to ∼3.5 mo, which allows investigation of the molecular programs underlying peak of neuronal diversity and amplification, older time points could reveal postnatal gene signatures that are being missed here. For example, astrocytes are prominent cell types that are shared among engineered and donor derived organoids and produce changes in gene expression and developmental trajectory. This finding could be further explored using older organoid samples, as astrocyte development is initiating at ∼day 100 and requires long term cultures to study their biology^92^. In addition, the developmental switch from *GRIN2B*-to *GRIN2A*-containing NMDARs occurs at ∼300 day old brain organoids^25^, which could potentially allow one to study postnatal human brain biology.

In the future, it will be important to expand upon this work by comparing this dataset with single cell transcriptomes obtained from *NRXN1* deletion carriers with other neuropsychiatric disorders like ASDs as well as those from healthy, unaffected individuals who also carry *NRXN1* deletions. This type of experimental design would allow dissection of the contribution of disease-specific effects at a greater scale – common vs. distinct molecular features across neuropsychiatric disorders which uniformly affect brain development and synaptic function. Furthermore, detection and analysis of alternative splicing of full-length transcripts at the single cell level across brain organoid development in these genetic contexts will better inform splicing misregulation. Finally, at the mechanistic level, why or how *NRXN1* deletions induce changes in alternative splicing will need to be determined. It is curious to know whether alternative splicing changes precede or is a response to synaptic dysfunction.

## Methods

### hPSC culture and forebrain organoid generation

hESC and iPSCs were cultured on feeder-free conditions as previously described^34, 35^. In order to form serum-free floating embryoid body (EB) aggregates, hPSCs were dissociated into single cells using Accutase (Innovative Cell Technologies). Dissociated cells were then reaggregated in low adhesion microwell culture plates (AggreWell-800, Stem cell technologies). 3 million cells were plated per 2mL well in mTesR plus Y-27632 (1 mM, Axon Medchem). 24 hours after plating, aggregated EBs were transferred to ultra-low attachment 10 cm petri dishes and cultured as previously described^89^. Briefly, EBs in 10 cm dishes are cultured in E6 medium (ThermoFisher) for up to 6 days with SB431542 (10 mM, Peprotech) and dorsomorphin (5 mM, Peprotech) for dual SMAD inhibition, promoting neural stem cell differentiation. Following day 6, EBs were cultured in Neurobasal (ThermoFisher) containing B27 without vitamin A supplement, Glutamax (Life Technologies), Penicillin-Streptomycin (ThermoFisher), Amphotericin B (Gibco) and the following morphogens at 20 ng/mL (Peprotech): human EGF, human FGF, human BDNF, and human NT3. At day 6-8, forebrain organoids were placed on an orbital shaker for gentle agitation to reduce spontaneous fusion. Starting at day 43, all morphogens were removed and brain organoids were cultured solely in B27 containing Neurobasal media.

### Lentivirus generation

Lentiviral plasmid constructs used in this study are Cre-recombinase and Flp-recombinase fused to EGFP driven by the ubiquitin-C promoter as previously described^35^. For all lentiviral vectors, viruses were produced in HEK293T cells (ATCC, VA) by co-transfection with three helper plasmids (3.25 μg of pRSV-REV, 8.1 μg of pMDLg/pRRE and 10 μg of lentiviral vector DNA per 75 cm^2^ culture area using calcium phosphate transfection method^90^. Lentiviruses were harvested from the medium 48 hrs after transfection. Viral supernatants were then centrifuged at a high speed of 49,000 x g for 90 min and aliquoted for storage in -80C. Viral preparations that yielded 90% EGFP expression were assessed to be efficiently infected and used for experiments.

### Cryopreservation and sectioning

Organoid samples were collected at day 21, 50 and 100. Samples were fixed in 4% paraformaldehyde at 4°C overnight then submerged in 30% sucrose/PBS solution for 24-48hrs in 4°C. Organoids were flash frozen in gelatin solution (gelatin in 10% sucrose/PBS) using dry ice/ethanol slurry and were stored in -80°C for long term storage or until cryosectioning. Cryosections were between 12 to 25 micron section thickness. Organoid sections were directly adhered to microscope slides and subsequently used for immunohistochemistry or stored for long term storage in -20°C.

### Immunostaining

Organoid sections were washed three times in 0.2% Triton-X in PBS (0.2%PBS/T) and then blocked in 10% normal goat serum diluted in 0.2%PBS/T (blocking solution) for 1hr at room temperature. Sections were incubated in primary antibodies diluted in blocking solution overnight at 4°C and were subsequently washed three times with 0.2%PBS/T, followed by incubation with secondary antibodies and DAPI diluted in PBS/T at room temperature for 2 hours. Finally, sections were washed three times (20 minutes per wash), and then mounted using Fluoromount mounting media (Southern Biotech). Primary antibodies used are as follows: mouse anti-Ki67 (1:250, BD Biosciences BDB550609), rabbit anti-SOX2 (1:500 Cell Signaling 3697S), rabbit anti-HOPX (1:500, Proteintech 11419-1-H), rat anti-CTIP2 (1:2000, Abcam ab18465), rabbit anti-TBR2 (1:1000, Abcam ab23345), mouse anti-SATB2 (1:1000 Abcam ab51502), mouse anti-NEUN (1:500, EMD Millipore MAB377), rabbit anti-NEUN (1:1000, EMD Milipore ABN78), rabbit anti-S100B (1:1000, Sigma S2644), chicken anti-MAP2 (1:5000, Abcam ab5392), rabbit anti-SYNAPTOPHYSIN (1:1000, Abcam ab14692), rabbit anti-HOMER (1:1000 Synaptic System 160003), and mouse anti-SYNAPSIN (1:500, Synaptic System 111011). Secondary antibodies conjugated with Alexa 488, 594, 647 (Invitrogen) and DAPI (1:1000, Sigma MBD0015) were used.

### Calcium imaging

Organoids were incubated in 1 mM of X-Rhod-1 AM dye (Invitrogen) diluted in a modified HEPES buffer (130mM NaCl, 5mM KCl, 2mM CaCl2, 1mM MgCl2, 10mM HEPES, 10mM Glucose, ∼pH 7.4 adjusted with NaOH) for 15 minutes at room temperature. Excess dye was washed with modified HEPES buffer once, then imaged using a confocal microscope (Nikon, A1R25). For GCaMP based calcium imaging, organoids were infected with pAAV-CAG-SomaGCAMP6F2 (Addgene) and imaged 2 weeks later. Imaging was carried out in glass bottom petri dishes (MatTek). Temperature was maintained at 37°C using the Ibidi stage heater. Time lapse images were acquired at 250 ms intervals for a period of 5 mins.

For picrotoxin stimulation, organoids were incubated for an hour in HEPES buffer containing picrotoxin (50 µM, Tocris Bioscience). Organoids were rinsed three times in drug free HEPES buffer and imaged. To inhibit activity stimulated by picrotoxin, organoids were placed in HEPES buffer containing CNQX/APV (2 µM/50 µM, Tocris Bioscience) and imaged immediately.

### Glycine stimulation

Organoids were washed once in Mg+ free HEPES buffer (130mM NaCl, 2.5mM KCl, 2mM CaCl2, 25mM HEPES, 30mM Glucose, ∼pH 7.4 adjusted with NaOH) and imaged for baseline activity using a confocal microscope (Nikon A1R25). Subsequently, organoids were incubated in Mg+ free HEPES buffer containing glycine (200 µM), strychnine (10 µM), and picrotoxin (50 µM) for 15 minutes. After incubation, organoids were rinsed three times then imaged for stimulated activity. Similarly for APV inhibition experiments, organoids were incubated in Mg+ free HEPES buffer containing glycine (200 µM), strychnine (10 µM, Fisher Scientific), and APV (50 µM) for 15 minutes, rinsed three times with drug free HEPES buffer and imaged, Temperature was maintained at 37°C using the Ibidi stage heater. Time lapse images were acquired at 250 ms intervals for a period of 5 mins for all experiments.

### Calcium imaging analysis

Images were processed using ImageJ software to produce binary images. Analysis was then carried out using a stimulation-free Matlab protocol as demonstrated previously^91^. Using the MATLAB protocol, we first ‘stacked’ the time lapse images captured by the confocal microscope to produce a Maximum Intensity Projection (MIP). Guided by the MIP, we then selected 4-5 regions of interest (ROI) indicating the most active regions of the organoid, with each ROI measuring 50 μm in diameter. Next, we selected the time interval which is determined by the recording duration (in seconds)/frames; in our recordings we used 300/109 for a time interval of 2.75. Changes in image intensity within ROI’s are then quantified and plotted as raw calcium traces. Using the average calcium intensity across all ROI’s in one field of view, synchronous spikes were plotted and a synchronous firing rate was determined using the number of detected synchronous spikes every minute. Frequency was determined by the total number of detected peaks every minute across all traces. Amplitude was established using the mean value of dF/F0 from individual peaks.

### Quantification and statistical analysis of calcium imaging data

Data wrangling was performed in Microsoft Excel, and all raw data points were transferred to Prism (9.3.0) for basic statistics, outlier detection, significance tests, and graph generation. To identify outliers from pooled replicates, the ROUT outlier test was used to identify outliers by fitting data with nonlinear regression and using a false discovery rate of Q=1%. To test for statistical significance, unpaired parametric two-tailed Student’s t-test was performed to compare the two genotypes and one-way ANOVA test with multiple comparisons (Tukey’s test) was used to compare conditions involving more than two samples.

### Live single cell dissociation

Organoids were rinsed 3 times with HBSS (10X HBS salt, 1M HEPES, 0.004M NaHCO3 diluted to 1X), then minced into small pieces and transferred to a 15mL conical tube for incubation in digestion solution (consisting of HBSS, 1 mg/mL Papain, 0.5mM EDTA, and 1mM L-cysteine) for 15 minutes at 37°C. Upon incubation, digestion mixture containing organoids were gently triturated with DNase I (25 μg/mL, Worthington-Biochem) and subsequently incubated again for another 10 minutes at 37°C followed by filtration with 70 µM and 30 µM filters (Miltenyi Biotech). Cell mixture was then centrifuged and pelleted. Cell pellet was resuspended in Neurobasal media with B27. This step was done to help dilute any remaining enzyme, EDTA, and other components of the digest mix. After the final re-suspension in Neurobasal media (without supplements), single cell mix was filtered again using 40 µM FlowMi pipet (Milipore Sigma) to help remove debris.

### 10XscRNASeq Protocol

Following isolation of single cells from brain organoids, cells were centrifuged at 300 g for 5 min and then re-suspended in 1 mL ice-cold Neurobasal media. Cell concentration and viability were determined using a hemocytometer with trypan blue dye exclusion and cell concentrations were adjusted to 700-1200 cells/µL for 10X single cell sequencing. For each sample, 9,600 cells were loaded into the 10X Chromium controller to target recovery of 6,000 cells and a Gel Beads in Emulsion (GEM) was generated. 10X Genomics 3’v3.1 chemistry was used. The samples were processed according to the protocol from 10X Genomics, using 14 cycles for cDNA amplification. Single cell libraries were sequenced using the Illumina NovaSeq 6000.

### Methanol fixation of brain organoids and processing of methanol fixed samples for 10X scRNASeq

Methanol fixation and rehydration of single cell suspensions were processed according to the 10X demonstrated protocol. Organoids were dissociated into single cells as described above. Dissociated cells were then centrifuged at 300 rcf for 5 mins at 4°C. Upon centrifugation, supernatant was removed and cells were triturated with 1mL of cold DPBS using a wide bore pipet tip. Cells were then centrifuged again at 300 rcf for 5 mins at 4°C and resuspended in 200uL of cold DPBS. This was immediately followed by adding 800uL of cold methanol dropwise to prevent cells from aggregating. Finally, the cell mixture was triturated gently to ensure it was well mixed. Methanol fixed cells were incubated in -20°C for 30 minutes and transferred to -80°C for long term storage.

The MeOH-fixed cells were retrieved from the -80 freezer, equilibrated on ice for 5 min, followed by centrifugation at 1,000 g for 5 min at 4°C. After centrifugation, supernatant was removed and the fixed cells were re-suspended in 500 uL wash/re-suspension buffer (0.04% BSA + 1 mM DTT + 0.2 U/uL Protector RNAse Inhibitor in 3X SSC buffer). The cells were visualized using trypan blue staining and concentration was adjusted to 700-1200 cells/uL for loading into the 10X instrument. 10X 3’v3.1 chemistry was used and 9,600 cells were loaded for each sample. The samples were processed according to the protocol from 10X Genomics, using 14 cycles for cDNA amplification. Single cell libraries were sequenced using the Illumina NovaSeq 6000.

### Single cell data alignment

10x single-cell RNA-sequencing data in Fastq files were aligned to transcripts using Cell Ranger 3.1.0 (https://www.10xgenomics.com/support/single-cell-gene-expression). Reference genome GRCh38 (Ensembl 93) was used as the reference genome. In the CellRanger *count* command, parameters *chemistry* and *expected-cells* were set as SC3Pv3 and 6000, respectively.

### Single cell preprocessing and normalization

Cell Ranger output h5 files were loaded using Seurat 4^92^ as the raw data. To reduce the impact of low-quality cells, we first removed cells with less than 1200 or more than 25,000 unique molecular identifiers (UMI). In addition, we removed cells with less than 600 or more than 6000 unique genes. Since low-quality or dying cells often exhibit extensive mitochondrial contamination, we removed cells with more than 10% mitochondrial transcripts.

Furthermore, we removed several clusters (details of the clustering will be mentioned later) of cells with low-sequencing depth to avoid the influence of poorly sequenced cells. Clusters with lower-than-normal distributions of the number of UMIs or unique genes were manually removed from the data. In the end, 9 lowly-sequenced clusters were removed from both donor-derived and engineered organoids single cell data (Figure S3). In addition, we utilized Scrublet^93^ investigate the doublets in the data. Only a small number of cells reached the threshold of doublets, indicating a low prevalence of doublets (Figure S4).

After the quality control, we finally harvested 33,538 genes and 156,966 high-quality cells, including 100,308 cells from 16 donor-derived organoid samples and 56,658 cells from 10 engineered organoid samples. Both original and processed data can be found in Data Availability. We normalized the total UMI counts per gene to 1 million (CPM) and applied log_2_(CPM+1) transformation for heatmap visualization and downstream differential gene expression analysis, which were conducted in Scanpy^94^. In the following Seurat integration procedure, we applied the default normalization approach of Seurat.

### Single cell integration

To reduce the influence of batch effects from multiple samples in single-cell data analysis, we applied the Seurat integration procedure to the data. We first loaded the raw data of each sample separately and created a list of Seurat objects after the quality control. Then we normalized each Seurat object and found the top 2,000 highly variable genes using the “vst” method in *FindVariableFeatures* function. 2000 integration features were selected from the series of Seurat objects using *SelectIntegrationFeatures*. Then integration features in each dataset were scaled and centered using the *ScaleData* function, based on which we ran the Principal Component Analysis (PCA) to reduce the high dimensions of features into 50 principal components.

In this study, we used the Reciprocal PCA (RPCA) procedure as the default method of integrating our large-scale data due to its high computational performance. We first identified integration anchors with previously identified integration features and top 30 reciprocal principal components, after which we ran the integration using the function IntegrateData with the top 30 dimensions for the anchor weight procedure. After integration, the data was scaled and PCA was conducted using *ScaleData* and *RunPCA* functions, respectively. Then the nearest neighbor graph was constructed using 20 k-nearest neighbors and 30 principal components. Louvain clustering was applied on the neighbor graph using the function *FindClusters* and multiple resolutions (0.5, 1.0, 2.0) were used to find clusters in both coarse and fine resolutions for comprehensive downstream analysis. Additionally, 2-dimensional and 3-dimensional embeddings of cells were generated using Uniform Manifold Approximation and Projection (UMAP) based on top 30 principal components. These integration, clustering, and dimensionality reduction procedures were applied to cells from donor-derived organoids, engineered organoids, as well as cells from both types of organoids.

In this study, we generated two batches of data and used reduced dimensions (principal components and UMAP embeddings) to analyze them. The first batch included 141,039 cells and 20 samples, on which we performed the dimensional reduction and cell type annotations. The second batch contained 6 samples. To integrate the embeddings and map the annotations of the new data onto the first batch of the data, we used the FindTransferAnchors and TransferData functions with the top 30 PC dimensions to learn anchors for transferring and transfer the data. We generated the integration PC embeddings using the IntegrateEmbeddings function and UMAP projections using the ProjectUMAP function. Finally, we predicted the cell type annotations of the cells in the new batch using the AddMetaData function.

### Cell Annotations

After the quality control and integration procedure, we got high-quality cells and clusters in multiple resolutions. A total of 49 clusters in a fine resolution (2.0) were generated from aforementioned procedures for cells from both donor-derived and engineered organoid samples. Canonical markers from previous studies were collected and used for manual annotations of each cluster, such as VIM for neural progenitor cells, STMN2 for neurons, and AQP4 for astrocytes. In addition, enrichment results of ToppCell-derived gene modules and prediction labels from reference datasets were used as supplementary evidence of annotations as well. A total of 29 cell classes were derived eventually, including subpopulations from NEC, glia cells, intermediate cells, neurons and supportive cells. Cluster 9 and 24 were labeled as unknown cells since there were no clear associations with known cell types based on marker genes or predicted cell types. Two clusters, including cluster 24 and 34, were labeled as low-sequencing-depth cells since their lower-than-normal transcript abundance levels. To focus on neuron differentiation, low-sequencing depth unknown cells, microglia cells, and mesenchymal cells were not included in the downstream analysis.

### Logistic regression for label prediction

To better understand the cell identities of clusters, we built up simple logistic regression models in the reference single cell data to predict cell type annotations in our own data. Such models were previously used in by Young et al.^95^ to infer the similarity between kidney tumor cell populations and known normal kidney cell types. In our study, we established logistic regression models as classifiers for each cell type in 4 public brain organoid single cell datasets and 1 fetal brain single cell dataset. The prediction scores from the models were used to classify whether one query single cell belongs to a specific cell type. We applied models of all cell types from reference data to each cell in our single cell data and calculated the average prediction scores of cell types or clusters. The results represent the association or similarity between reference and query cell types (Figure S6B).

### Differential expression analysis

In our study, we used the Wilcoxon test in the function *rank_genes_group* of Scanpy to calculate gene differential expression statistics. We applied the DE tests for comparisons between NRXN1 del cells and control cells in all cell classes and time points. Normalized expression values were used as the input data. FDR adjusted p values were used to control the type I error. Genes with FDR-adjusted p values lower than 0.05 in DE tests were defined as significant DE genes. In order to highlight DEGs relevant to our analysis (‘filtered’ list), we extracted and integrated a list of gene sets from Gene Ontology, including neurogenesis (GO:0022008), generation of neurons (GO:0048699), neuron differentiation (GO:0030182), neuron projection development (GO:0031175), neuron development (GO:0048666), neuron projection morphogenesis (GO:0048812), neuron projection (GO:0043005), somatodendritic compartment (GO:0036477), neuronal cell body (GO:0043025), myelin sheath (GO:0043209), axonal growth cone (GO:0044295). Additionally, we excluded genes associated with translational initiation (GO:0006413), ATP metabolic process (GO:0046034), and mitochondrion organization (GO:0007005). Both ‘filtered’ and ‘unfiltered’ DEG lists are shown in the Supplementary Tables. Volcano plots were generated for the visualization of DE genes using the *EnhancedVolcano* package^96^. In Figures 2 and 3, we conducted a hypergeometric test for each comparison of two gene lists to infer the significance of the number of overlapping genes in those two lists.- p value were corrected using FDR-adjusted p values.

### Gene modules from ToppCell

We used ToppCell toolkit to generate gene modules of cell types and clusters in our single cell data (Figure S6a)^97^. We applied ToppCell to user-provided cell annotations and derived well-organized gene modules for all cell classes. Each gene module contains the top 200 DEGs from ToppCell, representing the most prominent transcriptomic profile of this cell class. ToppCell-derived gene modules were seamlessly enriched using ToppGene^44^ and ToppCluster^98^.

### Gene enrichment analysis

Gene set enrichment analysis (GSEA) was conducted using ToppGene for gene sets from either ToppCell output or differential expression analysis. Gene ontologies were used to annotate molecular functions, biological processes and cellular components. In addition, we used the *prerank* function in GSEAPY package for the customized GSEA analysis. For UPS and alternative splicing enrichment gene plots (Figures S10 & S11), we consulted GSEA output from ToppGene as well as published literature reporting these genes. We used the manually curated neurological-disorder-associated gene sets^65^ as the reference for Figure 6a, such as genes of autism spectrum disorder and schizophrenia. We calculated FDR adjusted p values of enrichment for differentially expressed genes to infer their associations with neurological diseases. For disease enrichment analysis for GWAS loci (Table S10), we compiled a list of disease loci from the following publications: PMID: 28540026 (ASD)^64^, PMID: 31740837 (SCZ)^65^, PMID: 30718901 (MDD)^66^, and PMID: 31043756 (BP)^67^.

### Trajectory inference and pseudotime analysis

We used Monocle3^42^ to infer the pseudotime and trajectories of cell differentiations in the brain organoid single-cell data. We took advantage of the Seurat integration procedure and transferred Seurat objects into Monocle3 *cell_data_set* objects. Then we learned trajectories on the UMAP using the *learn_graph* function to get the pseudotime ordering of cells using the *order_cells* function. Cells with the highest expression levels of cell cycle genes in cycling NECs were selected as the start point of trajectories. In the end, every cell was assigned a pseudotime value, representing the estimated differentiation stages along the trajectory. Ridge plots were drawn based on the density of cells across pseudotime values.

### NRXN1 expression analysis

We collected human fetal cortex single-cell data from a large-scale single-cell dataset^40^. The normalized expression levels (log_2_(CPM+1)) of NRXN1 were calculated for each cell and average normalized expression was derived for cells in each cell types across various developmental stages.

### Alternative splicing and isoform analysis

We used bulk RNA-seq data from previous published induced neurons^31^. RNA-seq reads were aligned to the GRCh38.p13 reference genome using STAR version 2.7.9 using gene annotations from Gencode v41. Basic mode and “introMotif” were used for “twopassMode” and “outSAMstrandField” settings, respectively. Genes and isoforms were quanified with RSEM version 1.3.0. To measure the differential expression levels of genes and isoforms, we first loaded the output of RSEM with the package tximport, and then conducted differential gene expression and transcript usage analysis with the package DESeq2^48^. Differential transcript usage analysis was conducted for genotypes by controlling the effects of cell lines. FDR adjusted p values were used to control the type I error.

Majiq^50^ was used to quantify the alternative splicing. We first ran the Majiq Builder to de novo detect local splicing variations (LSVs) and build splicegraph. We then conducted Majiq Quantify to measure the Percent Splice In (PSI) and delta-PSI between genotypes. After quantification, we generated statistics of differential splicing variations and user interface for visualization using Volia. Additionally, we used ggsashimi^93^ to visualize the splicing events in sashimi plots with the bam file output from STAR.

### Reference Datasets

Several datasets were used for the cell type prediction in this study, including:

Kanton et al. (2019)^41^: This is a single cell dataset of human cerebral organoids derived from iPSC- and embryonic stem cell (ESC)-derived cells (43,498 cells) at different time points (day 0 ∼ day 120) during the differentiation.

Paulsen et al. (2022)^37^: This is a single cell dataset of human cerebral cortex organoids with haploinsufficiency in three autism spectrum disorder (ASD) risk genes in multiple cell lines from different donors of more than 745,000 cells.

Tanaka et al. (2020)^38^: This is synthetic analysis of single cell data from multiple brain organoid and fetal brain datasets. Data of 8 different protocols were collected and 190,022 cells were selected for the reannotation, where they classified 24 distinct clusters and 13 cell types.

Velasco et al. (2019)^24^: This is a study to validate the reproducibility of brain organoids with single cell sequencing. They collected 166,242 cells from 21 individual organoids and identified indistinguishable compendiums of cell types and similar developmental trajectories.

Zhong et al. (2018)^39^: This is a single cell dataset with more than 2,300 cells in developing human prefrontal cortex from gestational weeks 8 to 26.

Bhaduri et al. (2021)^40^: This is a large-scale single cell data of developing human brain from gestation week (GW) 14 to GW 25. Multiple brain regions and neocortical areas were sampled for the data.

## Data availability

The complete single cell-RNAseq dataset used for this study will be available upon publication.

## Code availability

Code used for single cell transcriptomic analysis can be found here (https://github.com/KANG-BIOINFO/NRXN1_effects). Calcium imaging analysis was performed in MATLAB using the previously published code (https://github.com/beccasbastian/CalciumIMG_Analysis).

## Acknowledgments

We thank Kelly Rangel (CCHMC gene expression core) and Dr. Jim Chambers (IALS Light Microscopy core) for assistance with 10X scRNAseq and Ca2+ imaging set up as well as members of the Pak lab for experimental assistance and helpful discussions. We also thank Dr. Zhiping Pang for sharing the psychiatric risk summary gene list for disease enrichment analysis.

## Author contributions

R.S., R.B., Y.J.S., and J.B. cultured brain organoids and performed experiments. K.J. carried out all scRNAseq data analysis. R.S. and N.P. conducted live Ca2+ imaging and analysis. A.P. and R.B. optimized single cell dissociation protocol. R.G. optimized image analysis. R.S., K.J., Y.B.S., B.A., and C.P. designed the experiments and R.S., K.J., and C.P. wrote the manuscript.

